# Population genetics of sugar kelp in the Northwest Atlantic region using genome-wide markers

**DOI:** 10.1101/2020.04.21.050930

**Authors:** Xiaowei Mao, Simona Augyte, Mao Huang, Matthew P. Hare, David Bailey, Schery Umanzor, Michael Marty-Rivera, Kelly R Robbins, Charles Yarish, Scott Lindell, Jean-Luc Jannink

## Abstract

An assessment of genetic diversity of marine populations is critical not only for the understanding and preservation of natural biodiversity but also for its economic potential. As commercial demand rises for marine resources, it is critical to generate baseline information for monitoring wild populations. Furthermore, anthropogenic stressors on the coastal environment, such as warming sea surface temperatures and overharvesting of wild populations, are leading to the destruction of keystone marine species such as kelps. In this study, we conducted a fine-scale genetic analysis using genome-wide high-density markers on Northwest Atlantic sugar kelp species, *Saccharina latissima* and putative species, *Saccharina angustissima*. The population structure for a total of 149 samples from the Gulf of Maine (GOM) and Southern New England (SNE) was investigated using AMOVA, Fst, admixture, and PCoA. Genome-wide association analyses were conducted for six morphological traits, and the extended Lewontin and Krakauer (FLK) test was used to detect selection signatures. Our results indicate that the GOM region is moderately more heterogeneous than SNE. While admixture was observed between regions, these results confirm that Cape Cod acts as a biogeographic barrier for sugar kelp gene flow. We detected one significant SNP (P-value=2.03×10^−7^) associated with stipe length, and 243 SNPs with higher-than-neutral differentiation. The findings of this study provide fundamental knowledge on sugar kelp population genetics for future monitoring, managing and potentially restoring wild populations, as well as assisting in selective breeding to improve desirable traits for cultivation and bioenergy production.

## Introduction

Brown macroalgae in the order Laminariales (Phaeophyceae), or kelp, are keystone species in the near-shore temperate marine environment. As primary producers, they are at the base of many food webs and provide numerous ecosystem functions including detrital production, wave attenuation, habitat modification, and carbon sequestration (Dayton 1985; Bartsch et al., 2008; Falkenberg et al., 2012; Efird and Konar 2014; Trebilco et al., 2015). For humans around the globe, kelp has long provided both a food source and bioextracts with various applications (Bartch et al., 2008).

Besides their important ecological roles and the well-established kelp cultivation practices in coastal Asian countries, there is growing interest in macroalgal cultivation in Europe, South America, and throughout the USA (Augyte et al., 2017; Buschmann et al., 2017; Campbell et al., 2019; Grebe et al., 2019; Kim et al., 2019; Geocke et al., 2020). Specifically, there are efforts to selectively breed kelp for large-scale food and bioenergy production (Bjerregaard et al., 2016; Hwang et al., 2019; Geocke et al., 2020) and increasing demand for germplasm banking to support future cultivation as well as restoration research (Barrento et al., 2016; Wade et al., 2020). To assist in the establishment of this nascent industry, an understanding of genetic variation across wild kelp populations is essential. Intensive selection pressure during the marine crop domestication process leads to favoring certain phenotypic traits (Zhang et al., 2017). These mechanisms may promote adaptive divergence between cultivated seaweeds and wild populations and it is, therefore, critical to have an understanding of wild phenotypic traits as they undergo domestication. This will foster future breeding and cultivation efforts in a sustainable and informed manner for managers, conservation groups, researchers, and industry.

Although kelp are key both ecologically and economically worldwide, the evolutionary history of kelp species is not entirely known (Bolton 2010; Starko et al., 2019). Limited research exists on the genetic and phenotypic structure of kelp within species, despite broad range distributions and likely regional adaptation (Valero et al., 2011). Existing *S. latissima* populations colonized the eastern and western North Atlantic coasts post-glacially from a north Pacific source via the oceanographic flow through the Arctic (Neiva et al., 2018). New evidence suggests that kelp have radiated at a constant rate, but members of the clade containing *Saccharina* show an additional rate increase (Starko et al. 2019).

Genetic population structure depends on the mode of reproduction and dispersal ability (Valero et al., 2011), and therefore provides insights about gene flow among populations (Durrant et al., 2018). In the marine environment, where direct observation of dispersal can be challenging, genetic tools provide an opportunity to better recognize patterns and scales of population connectivity (Valero et al., 2011). Nearshore kelp cultivation efforts have raised concerns by resource managers of the potential of farm cultivars to introgress with wild populations. However, for kelp species like *S. latissima*, dispersal distances of meiospores and gametes are generally short, usually not exceeding a few meters (Paine 1979; Hoffman and Santelices 1991). This was demonstrated by Zhang et al. (2017) where genetic structure and relatedness were evaluated in 8 wild populations and 17 farmed *Saccharina japonica* and they observed that wild populations have not been significantly impacted by gene flow from cultivated populations.

In the Northwest Atlantic, several small-scale studies have investigated the genetic population structure of *S. latissima* within small geographic ranges. A previous study in the Gulf of Maine, (GOM) reported slight genetic differentiation between five *S. latissima* populations spanning 225 km (Breton et al., 2018). Augyte et al., (2017) also found a slight distinction between the sugar kelp population at one site in Long Island Sound (LIS) as compared to three populations tested in the GOM. However, these and other population-level genetic studies of kelp may have been limited by variation levels in amplified fragment length polymorphisms (Vos et al., 1995) and microsatellites (Neiva et al., 2018; Richard et al., 2008; Nielsen et al., 2016; Paulino et al., 2016). Improvements in the speed, cost, and accuracy of next-generation sequencing (NGS) data can now provide better resolution to kelp genetics studies. In addition to NGS, reduced representational sequencing such as Genotyping by Sequencing (GBS) (Elshire et al., 2011) and Diversity Arrays Technology (DArT) (Jaccoud et al., 2001) have additional advantages, including reducing the genome complexity, avoiding inherent ascertainment bias in fixed SNP arrays, and decreasing sequencing costs. The high density of SNPs from these methods provides a finer understanding of the evolutionary processes shaping genetic diversity and, in some cases may be informative about genes affecting complex traits in wild populations.

Our study focused on *Saccharina latissima*, sugar kelp, which has a circumboreal distribution. In the Northwest Atlantic, its southern distributional limit is in Long Island Sound (LIS), with one disjunct historic population at an offshore site in New Jersey (Egan and Yarish 1988). Long Island Sound (41° N, 72-73° W) has a coastline of roughly 176 km and ranges from New York City in the west to Fisher’s Island in the east (Mathieson and Dawes 1988). To the north of LIS, separated by Cape Cod, is the Gulf of Maine (GOM) where a dominant coastal current flows southwestward (Pappalardo et al., 2015). This direction in water flow may contribute to sugar kelp gene flow moving southward via meiospore or rafting transport. The GOM is also home to an endemic kelp, *Saccharina angustissima*, closely related to *S. latissima* but with a distinctive strap-like morphology, found only within an eight nautical mile radius on intertidal ledges and islands exposed to heavy surf (Mathieson et al., 2008; Augyte et al., 2017; 2018).

The objectives of this study were twofold: 1) to explore the finer population structure of sugar kelp (*S. latissima* and *S. angustissima*) in the Northwest Atlantic and the admixture between regions separated by Cape Cod and 2) to test for associations between genetic variation and phenotypic trait variation. This study is fundamental to understanding the genetic diversity of sugar kelp in the context of species adaptation to environmental stressors, especially in the face of changing climate and warming oceans (Reusch et al., 2005). Furthermore, our work has the potential to inform recommendations for protecting coastal marine ecosystems and guide future kelp breeding and cultivation efforts by building a baseline of knowledge about kelp population diversity and connectivity.

## Materials and Methods

### Sample collection and phenotypic analysis

A total of 189 wild kelp samples were collected by SCUBA diving at 15 locations (Fig. 1) throughout the Northwest Atlantic conducted during April – June of 2018. Among these samples, we recorded six morphological traits on 165 samples from 15 locations (Table 1). Of these, care was taken to identify and only use reproductive blades, and thus it was assumed that most of the blades were at full maturity. The six traits were blade length, blade width at 10cm, blade width at the widest portion of the blade, blade thickness, stipe length, and stipe diameter. Collection locations were selected based on existing kelp beds and the ease of access (Table 1). Among these 15 locations, *S. angustissima* was found only at one location (Giant’s Staircase). The population sampled (defined as all 15 locations) was divided into two regions separated by Cape Cod, with GOM to the north of Cape Cod, and Southern New England (SNE) to the south. Radar plots of the mean and standard deviation of morphological traits were made to visualize diversity. Furthermore, pairwise correlations of the six traits were estimated using the cor() function in R (method = “pearson”, option = “na.or.complete”). The correlation was considered statistically significant when P-value < 0.05.

**Table 1.**
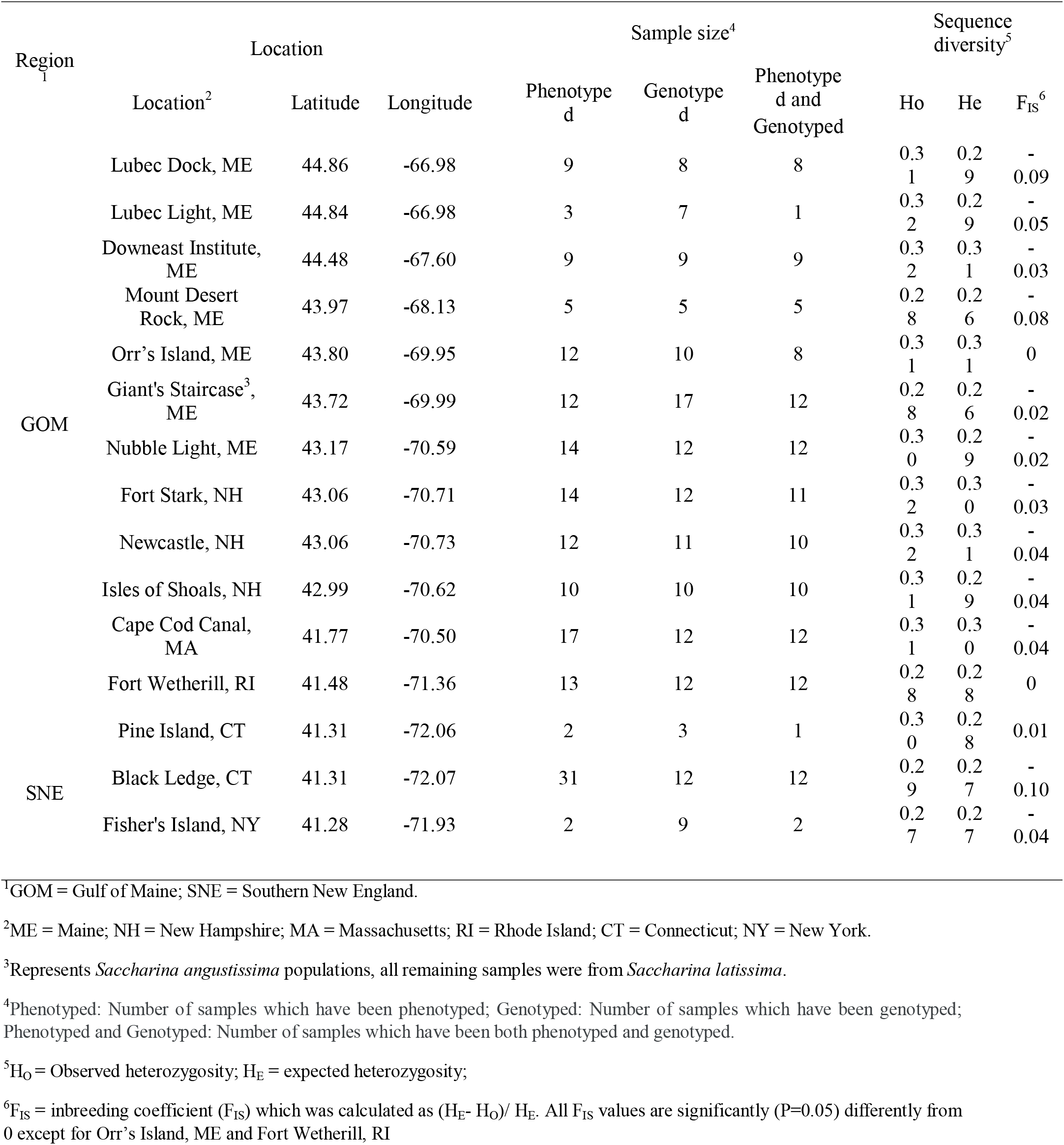
Location, sample size and sequence diversity for sampling locations.

**Figure 1.**
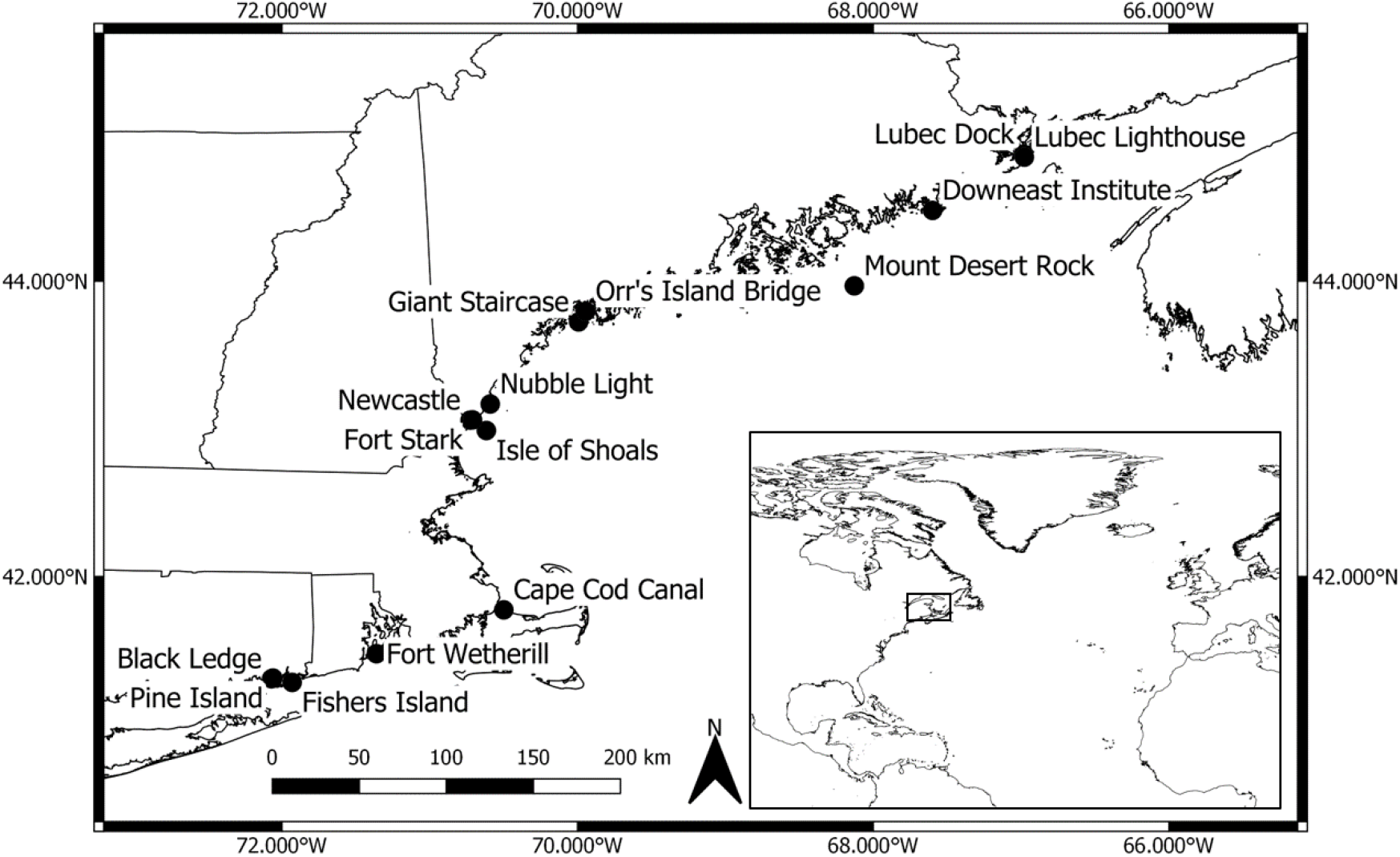
Collection locations of 15 subtidal populations of *Saccharina latissima* and *Saccharina angustissima* used in this study.

### DNA extraction and genotyping

Of the 189 collected samples, 149 kelp samples from both GOM and SNE for 15 locations were genotyped ￼(Table 1). ￼Meristematie kelp blade tissue was removed and placed in silica gel to desiccate. After transport to UCONN Stamford, Marine Biotechnology Laboratory, samples were ground up with mortar and pestle using liquid nitrogen. Weights of 5 mg per sample were measured and sent to DArT Ltd (Diversity Arrays Technology, Canberra, Australia) sequencing facility for DNA isolation and genotyping. Genomic DNA was extracted according to the modified CTAB protocol (Doyle and Doyle, 1990). Then DNA quality and quantity were evaluated with a DS-11FX series spectrophotometer (Denovix, Wilmington, DE, USA), followed by running agarose (1.2%) gel electrophoresis. Sugar kelp genotyping was carried out in two steps: First, reduced genomic representations were generated following the procedures described by Kilian et al (2012). This procedure included the digestion of DNA samples using a rare cutting enzyme PstI and secondary frequently cutting restriction endonucleases (RE), ligation with site-specific adapters, and amplification of adapter-ligated fragments. Second, nextgeneration sequencing technology was applied to detect SNPs and silicoDArT markers using HiSeq2000 (Illumina, USA). The sequence data were analyzed using DarTsoft14 (an automated genotypic data analysis program) and DArTdb (a laboratory management system). Two technical replicates of each DNA sample were genotyped to guarantee the reproducibility of the sequencing. The detailed procedure can be found in Kilian et al., (2012).

### Quality control for SNPs

A total of 20,242 SNPs were identified and five steps of quality control were applied to the SNP data: 1. Removal of sequence tags with more than one SNP; 2. Removal of SNPs with call rate (proportion with non-missing samples) less than 95%; 3. Removal of samples with call rate (proportion with non-missing SNPs) less than 90%; 4. Removal of SNPs with minor allele frequency (MAF) < 0.05; 5. Removal of SNPs severely departing from Hardy-Weinberg-Equilibrium (P-value < 0.01) in more than ¼ of all collection sites. After the quality control, a total of 149 samples and 4,905 SNPs were retained.

### Sequence diversity and population structure

The sequence diversity for samples within each location was assessed by summary statistics of average expected heterozygosity (H_E_), average observed heterozygosity (H_O_), and inbreeding coefficient (F_IS_), using the R package *hierfstat* (Yang 2006).

Population structure was assessed through analysis of molecular variance (AMOVA), pairwise F_ST_, Principle Coordinate Analysis (PCoA) and admixture analysis. First, the presence of a hierarchical population structure was investigated using the AMOVA (Excoffier et al., 1992) implemented in the R package *poppr* (Kamvar et al., 2015). In AMOVA, hierarchical variance components of GOM and SNE were estimated both combined and separately, where *S. angustissima* and *S. latissima* were analyzed together, due to their similarities inferred from PCoA. Second, pairwise relationships among all locations were investigated by calculating pairwise F_ST_ using the R package StAMPP (Pembelton et al., 2013). The AMOVA and pairwise F_ST_ rely on geographical assignments, estimating genetic differentiation among different locations. The underlying population structure was further investigated through PCoA implemented in the R package *Adegenet* (Jombart 2008). The PCoA is a model-free approach, which is commonly used to find hidden structure among samples as the population is not preassigned. Admixture analysis was done in R using the conStruct package (Bradburd et al., 2018). Similar to other population structure analyses, this model assumes a number K of distinct ancestral populations, called “layers”. Distinct from other models, conStruct fits isolation by distance (IBD) within layers. Bradburd et al. (2018) showed this model better captures the true subpopulation structure when compared to conventional non-spatial admixture modeling. Models with different values of K are fit and a “best model” can be chosen either based on its crossvalidation prediction of covariances in allelic frequencies across samples or based on the contribution of layers to explaining allelic frequency covariance. Due to computational intensity in conducting the cross-validation approach, we used the latter approach. The model was initially fit using values of K=1 to K=10 for the full set of samples and K=1 to K=6 for the GOM samples using a smaller number of iterations (8,000). Based on these results, we reduced the values of K to 1 to 6 for the full set and 1 to 4 for the GOM samples using a larger number of iterations (50,000 MCMC iterations with a burn-in of 500 iterations that were discarded). The final selection of K was chosen based on: the minimum contribution for a layer to be included was 1% of the total covariance (Supplemental Fig. 1). This resulted in K=3 for all samples tested, and K=2 for just the GOM samples.

### Isolation by distance

The extent of IBD (Wright 1931), which assumes that populations that are geographically closer tend to be more closely related genetically, was also estimated for our samples using a Mantel test implemented in R package *VEGAN*. In the Mantel test, genetic distance was represented by F_ST_/(1-F_ST_), and geographical distance was calculated as “the crow flies” (straight line) between our collection sites. The Mantel test was run for GOM and SNE separately with 999 permutations to assess the significance of the correlation between genetic and geographic distance.

### Genome-wide association analyses of morphological traits

A genome-wide association study (GW AS) was conducted on six traits using the 4,905 SNP markers and 98 GOM plus 27 SNE samples (a common set out of the 149 genotyped and 165 phenotyped samples). Due to the strong subpopulation structure in this data set and to account for sampling location variation, the sampling location (15 locations) was included as a categorical fixed effect in the GWAS model using the R package *GAPIT* (Lipka et al., 2012). The kinship matrix was estimated using *rrBLUP* package (Endelman, 2011) with the A.mat() function and included in the GWAS model. The mixed linear model was as follows:

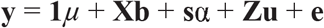

where **y** was a vector of the response variable for one of the morphological traits, **1** was a vector of ones, *μ* was the common population mean, **b** was the vector of location effects, **X** was an incidence matrix relating location to each individual, α was the additive allele substitution effect of the SNP, and **s** was the design matrix for SNPs. Elements of the vector **s** were allele dosages (0 to 2) for one randomly chosen allele at each locus. **Z** was an incidence matrix relating the vector of additive polygenic values **u** to individuals, and **e** was the error term. The values **u** and **e** were assumed to follow multivariate normal distributions, 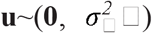 and 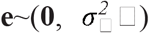 respectively, where ***G*** was the genomic covariance matrix estimated by *GAPIT* using the kinship matrix. ***I*** was an identity matrix, 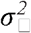 was the genetic variance, and 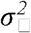 was the residual variance. To correct for the multiple testing, a Bonferroni correction was used. The resulting significance level threshold was 0.05/4,905 = 1.02 × 10^−5^. SNPs with a P-value below this threshold were significantly associated with morphological traits.

### Detection of selection signatures

The FLK test (Bonhomme et al., 2010) was used to test for signatures of selection. This test can identify SNPs with especially high differentiation among populations, so-called outliers relative to the neutral background, while accounting for the hierarchical structure among populations. We applied the FLK test to 149 genotyped samples from 15 locations which included both GOM and SNE samples where GOM included both *S. latissima* and closely related *S. angustissima*. There were five steps for running the FLK analysis with detailed information described by Bonhomme et al. (2010): 1. Compute Reynolds’ genetic distance from SNP data; 2. Build rooted Neighborjoining tree using the R package *APE* (Paradis et al., 2004); 3. Compute population kinship matrix from Neighbor-joining tree; 4. Compute FLK test statistic; 5. Simulate empirical distribution of FLK test statistic under the null hypothesis of neutral evolution with 50,000 replicate and return the empirical quantiles of the null distribution. FLK test statistics above 0.995 quantiles were considered to be significant. This procedure was applied to the 4,905 SNPs for all populations.

## Results

### Phenotypes of collected samples

Large morphological variation was observed across locations for the collected samples (Fig. 2). Blade lengths ranged from 84.5 ± 37.5 cm for Pine Island to 227.4 ± 22.9 cm for Mount Desert Rock; blade widths at 10 cm ranged from 2.2 ± 0.3 cm for Giant Staircase to 24.6 ± 1.6 cm for Downeast Institute; blade widths at the widest ranged from 3.4 ± 0.3 mm for Giant Staircase to 41.4 ± 2.3 mm for Orr’s Island; stipe diameter ranged from 2.17 ± 0.61 mm for Giant Staircase to 14.43 ± 2.17 mm for Downeast Institute; stipe lengths ranged from 4.8 ± 0.6 cm for Giant Staircase to 122.7 ± 18.9 cm for Lubec Light; blade thickness ranged from 0.8 ± 0.00 mm for Pine Island to 2.28±0.14 mm for Downeast Institute. Most of the pairwise correlations were positive except for the correlation between blade width at 10cm and blade length which was - 0.05 but not significantly different from zero (Fig. 3). Among all positive correlations, stipe length was most highly correlated with stipe diameter (0.85), and blade width at the widest portion was most highly correlated with blade width at 10 cm (0.80).

**Figure 2.**
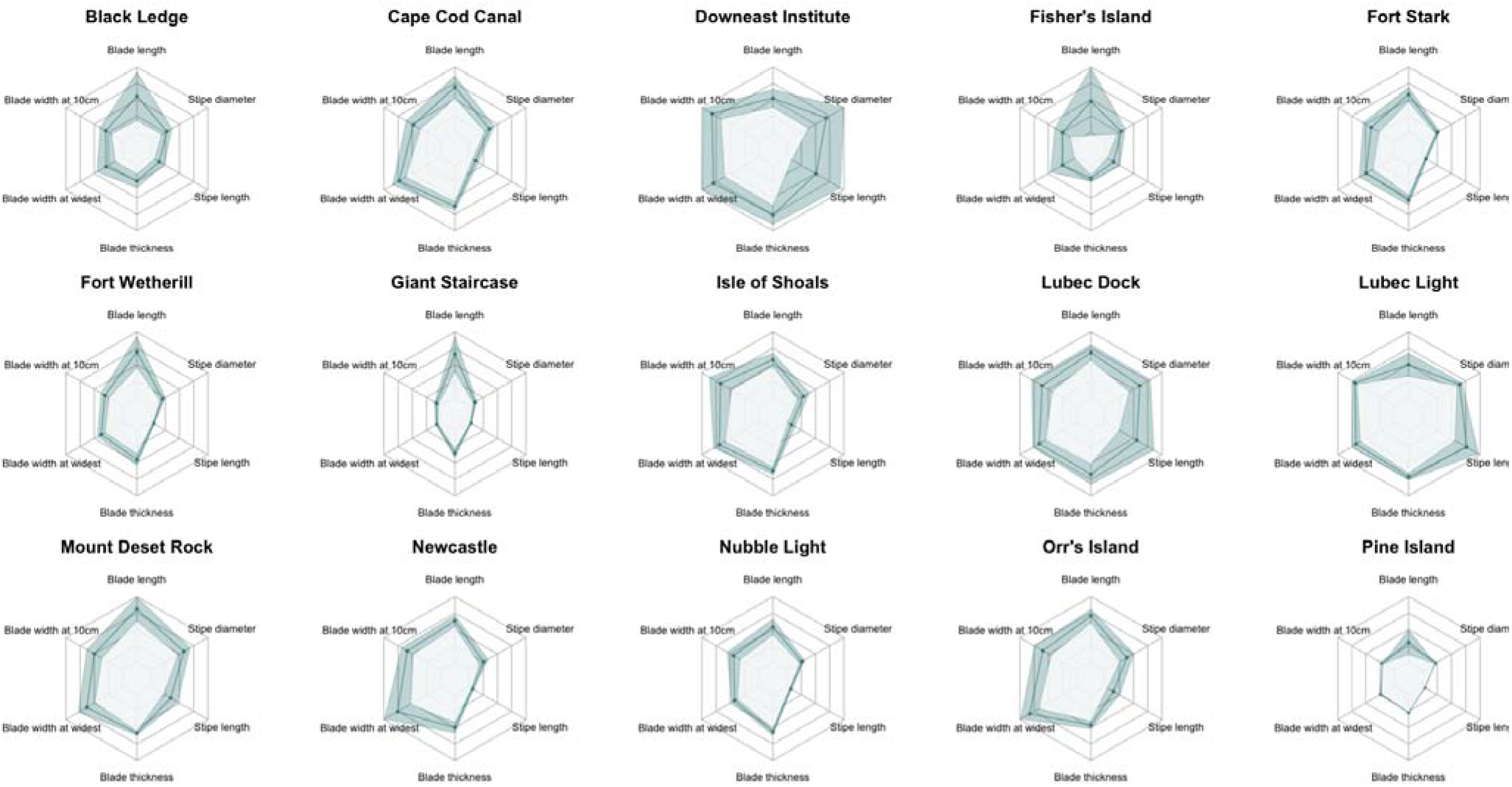
The radar plots for the phenotypic mean ± standard error (darker shading) in 15 sampling locations. The maximum value for the axis for each trait were as follows; 280.0 cm for blade length, 30.0 cm for blade width at 10 cm, 50.0 cm blade width at widest, 3.0 mm for blade thickness, 160.0 cm for stipe length, and 21 mm for stipe diameter.

**Figure 3.**
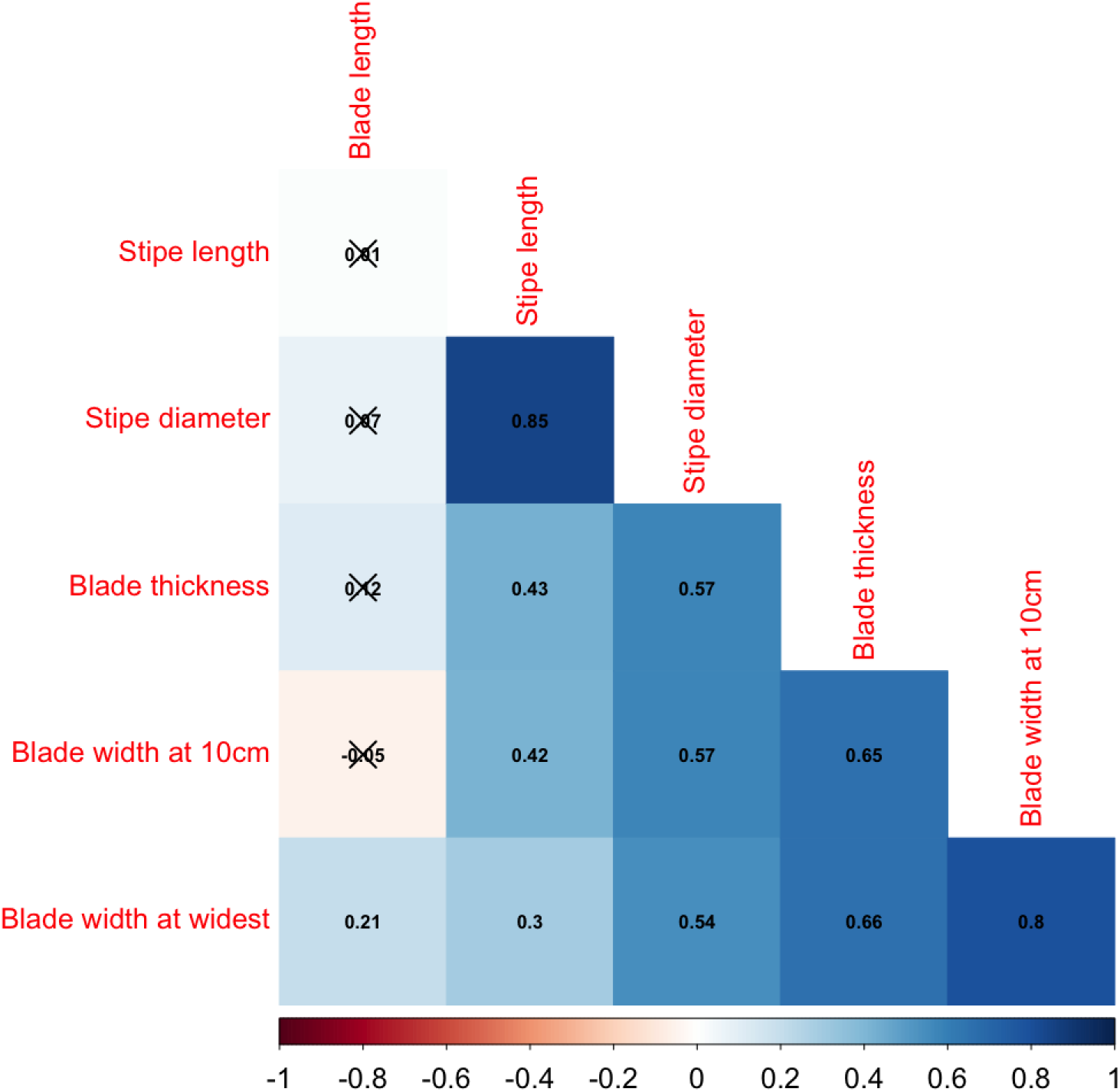
Phenotypic correlations among six morphological traits. The color gradient is shown at the bottom with red representing a negative correlation and blue representing a positive correlation. Non-significant (P-value >0.05) correlations were marked with black crosses.

### Sequence diversity

Most locations had a slightly negative inbreeding coefficient (F_IS_) (Table 1). Observed heterozygosity (Ho) values ranged from 0.27 for Giant’s Staircase to 0.322 for Newcastle. Overall, expected and observed heterozygosity was similar, suggesting random mating in all sampled locations. Lubec Dock had the most negative F_IS_ and Pine Island had the most positive F_IS_ among all locations.

### Population structure

The AMOVA results indicate that roughly half of the total variation exists within locations (56.4%), while the least variation exists among locations within each region (14.4%, Table 2), similar to what has been reported by Geocke et al., (2020). Variation among locations was higher in GOM (23.0% of the total variance) than among locations in SNE (6.8%, Table 2). Populations in SNE were more genetically homogenous than populations in GOM: the average pairwise F_ST_ among SNE locations was 0.03, as compared to 0.13 among GOM locations. More revealing was that pairwise F_ST_ showed that GOM was distantly related to SNE (F_ST_ > 0.25, Fig. 4, Supplemental Table 1). The PCoA revealed two major clusters, dividing samples into GOM and SNE regions (Fig. 5). The admixture analysis using conStruct revealed three underlying ancestral populations for all our 149 samples (Fig. 6). There was a clear difference between GOM and SNE, corresponding well to the pairwise F_ST_ analysis and supported by the PCoA results (Fig. 5). Yet samples from SNE share some ancestry with samples from GOM.

**Table 2.** Table 2. Analysis of molecular variance for sugar kelp in Northwest Atlantic using genome-wide single nucleotide polymorphisms data.

| Scenario | Source of variation | Variance | Percentage (%) |
| --- | --- | --- | --- |
| All locations | Between regions | 349.55 | 29.26 |
|  | Among locations within region | 171.97 | 14.39 |
|  | Within location | 673.25 | 56.35 |
| GOM<sup>1</sup> | Among locations within region | 202.62 | 23.01 |
|  | Within location | 678.11 | 76.99 |
| SNE<sup>2</sup> | Among locations within region | 47.98 | 6.80 |
|  | Within location | 657.80 | 93.20 |

<sup>1</sup>GOM = Gulf of Maine;
<sup>2</sup>SNE = Southern New England.

**Figure 4.**
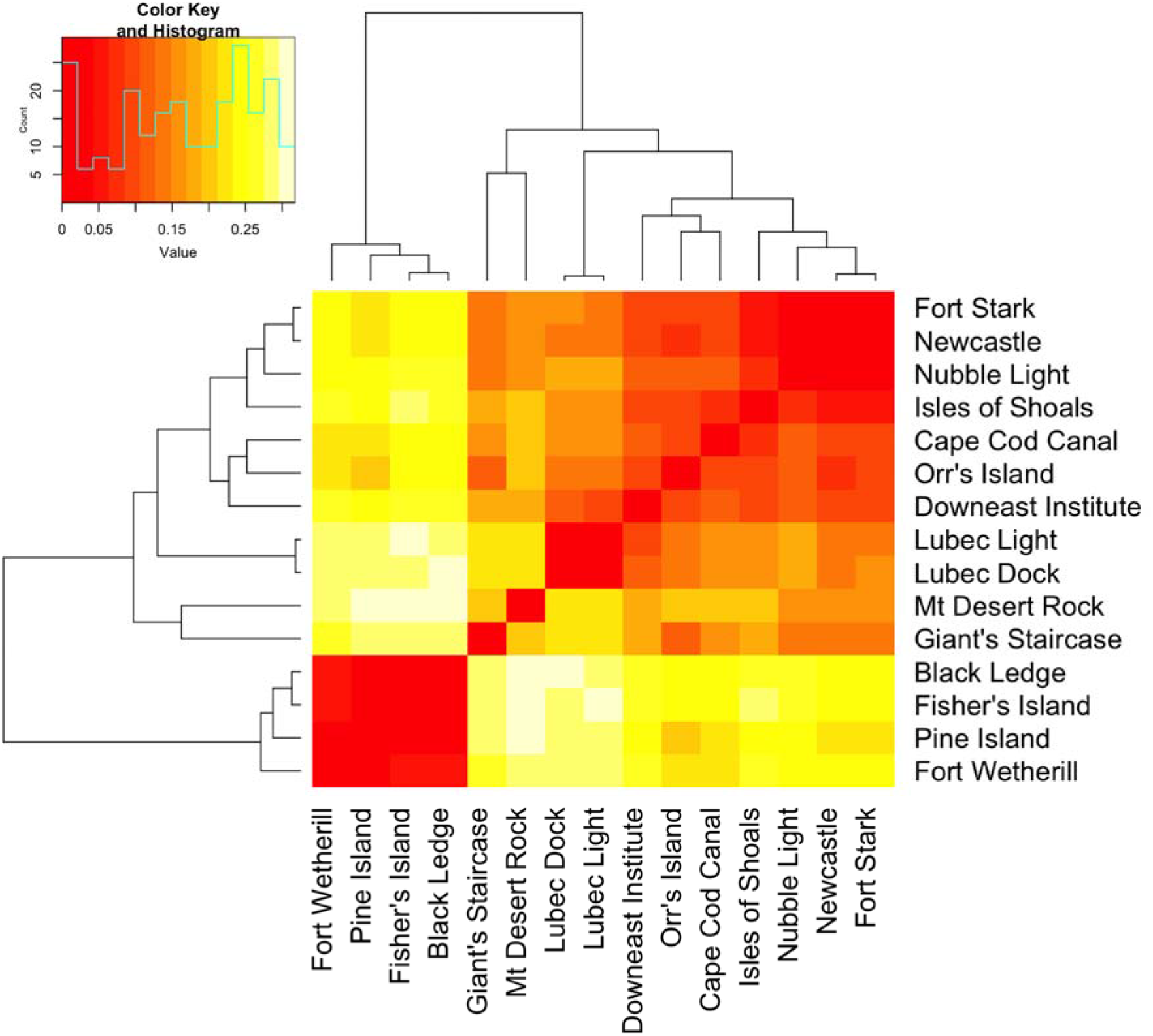
Heat map showing pairwise F_ST_ values with hierarchical clustering. Insert graph on the top left is a histogram showing the frequency spectrum of pairwise F_ST_ values.

**Figure 5.**
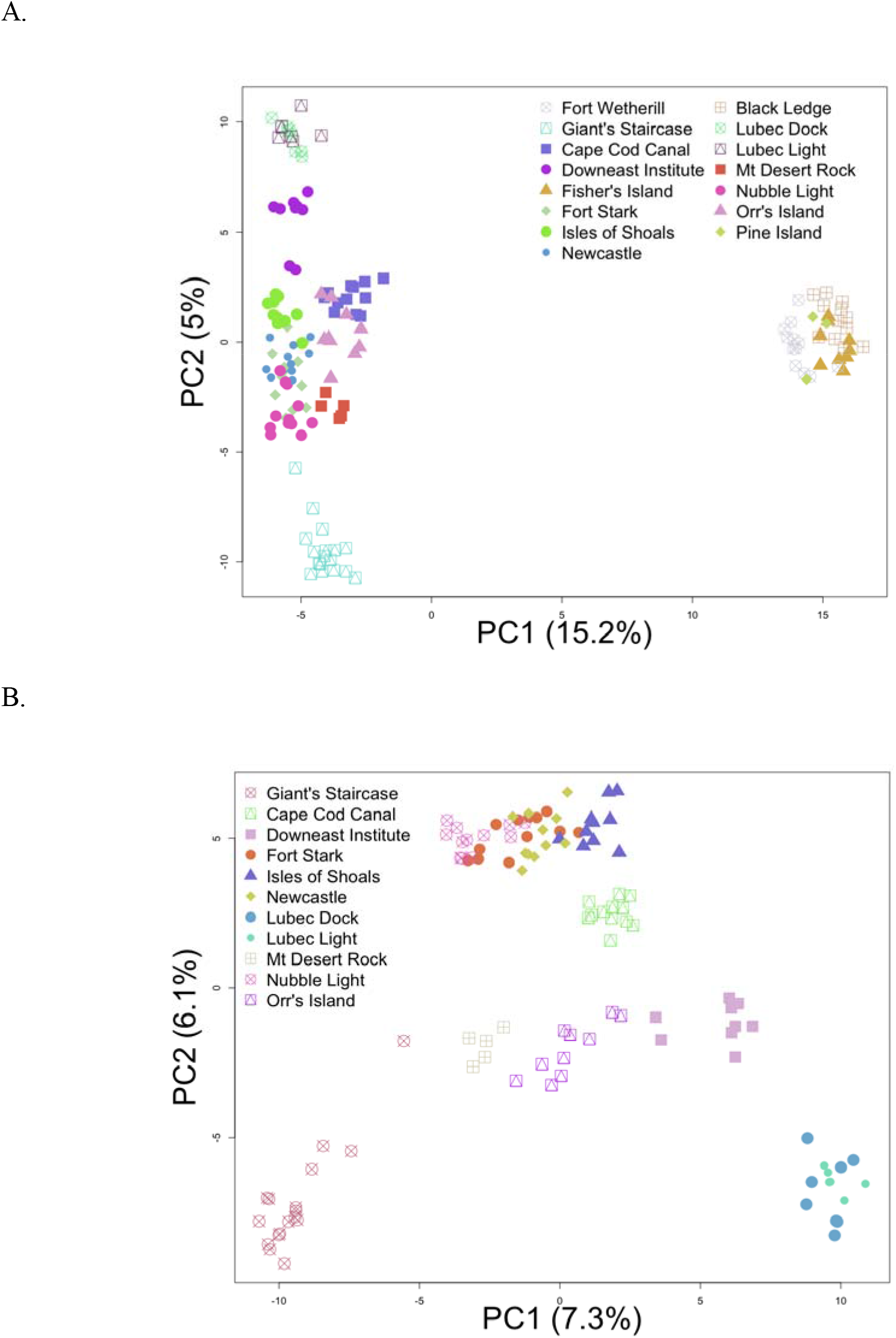
Principal coordinate analysis (PCoA). Samples are labeled according to their collection locations. The two PCs that explain most of the variation are shown in the plots. (A.) PCoA of 149 samples from all 15 locations; (B.) PCoA of 113 samples from GOM (Gulf of Maine) only.

**Figure 6.**
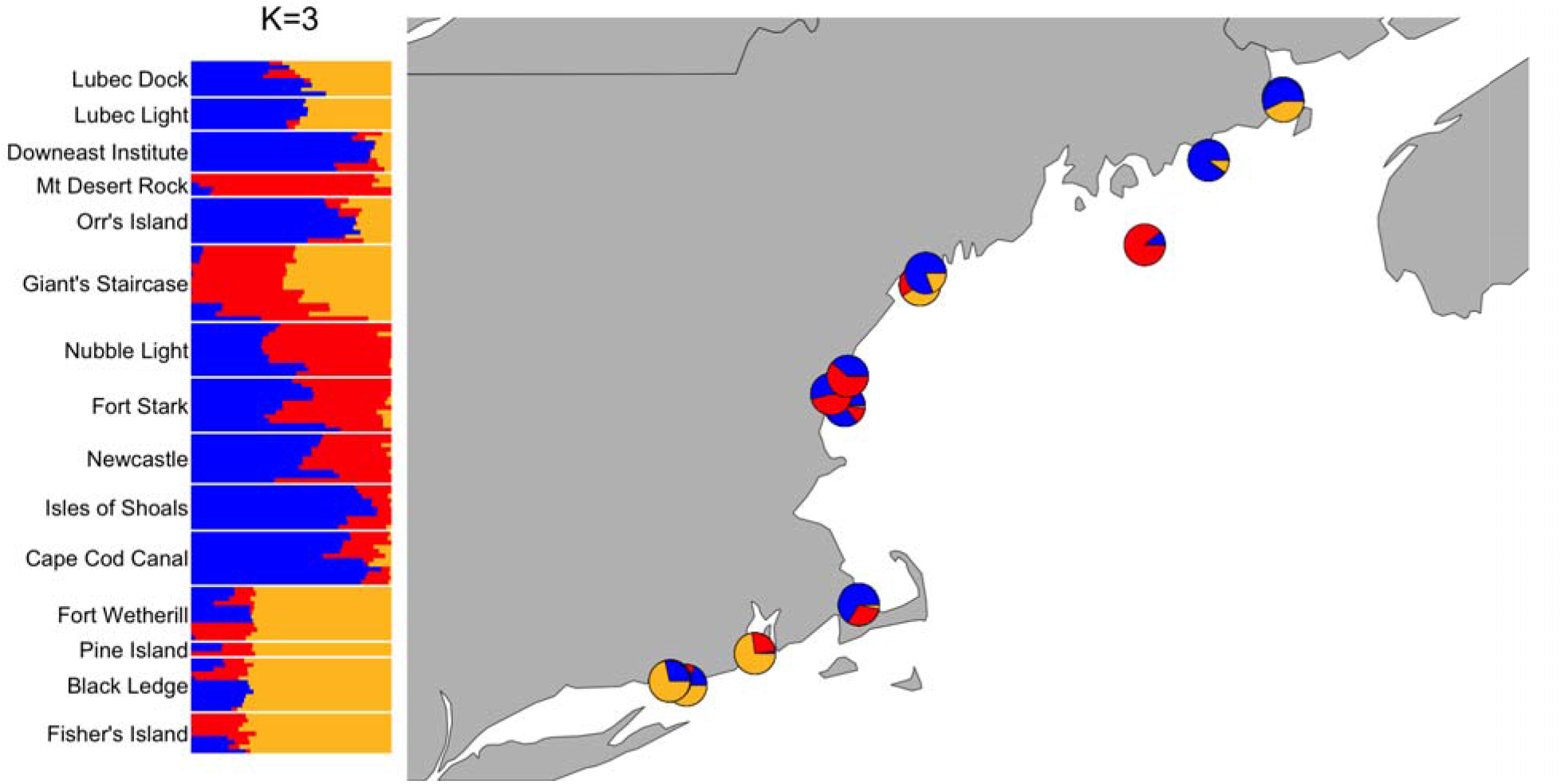
Admixture proportions estimated for the 149 kelp samples for K=5 (left). Admixture proportions are shown as pie plot at their approximate geographic locations (right).

### Isolation by distance

Testing IBD for each region separately, the Mantel tests (Fig. 7) showed a moderately positive correlation for GOM (r = 0.47, P-value = 0.002) and a strong positive correlation for SNE (r = 0.94, P-value = 0.125). The strong IBD was not significant for SNE due to the small number of populations (n = 4).

**Figure 7.**
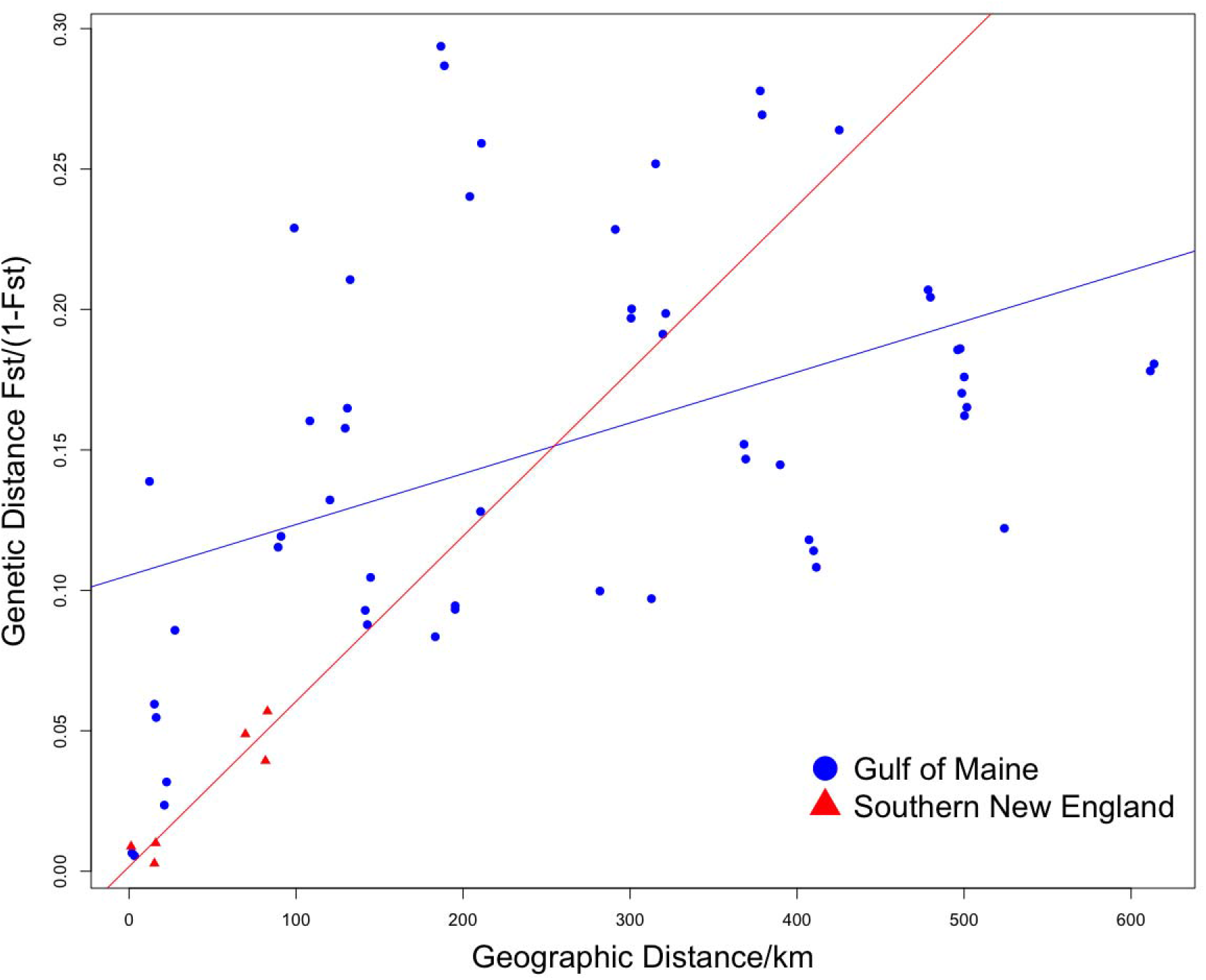
Mantel tests for correlation between genetic distance (y-axis) and geographic distance in kilometers (x-axis) in the Gulf of Maine and Southern New England. Regression lines are drawn for the Gulf of Maine (blue) and Southern New England (red).

**Figure 8.**
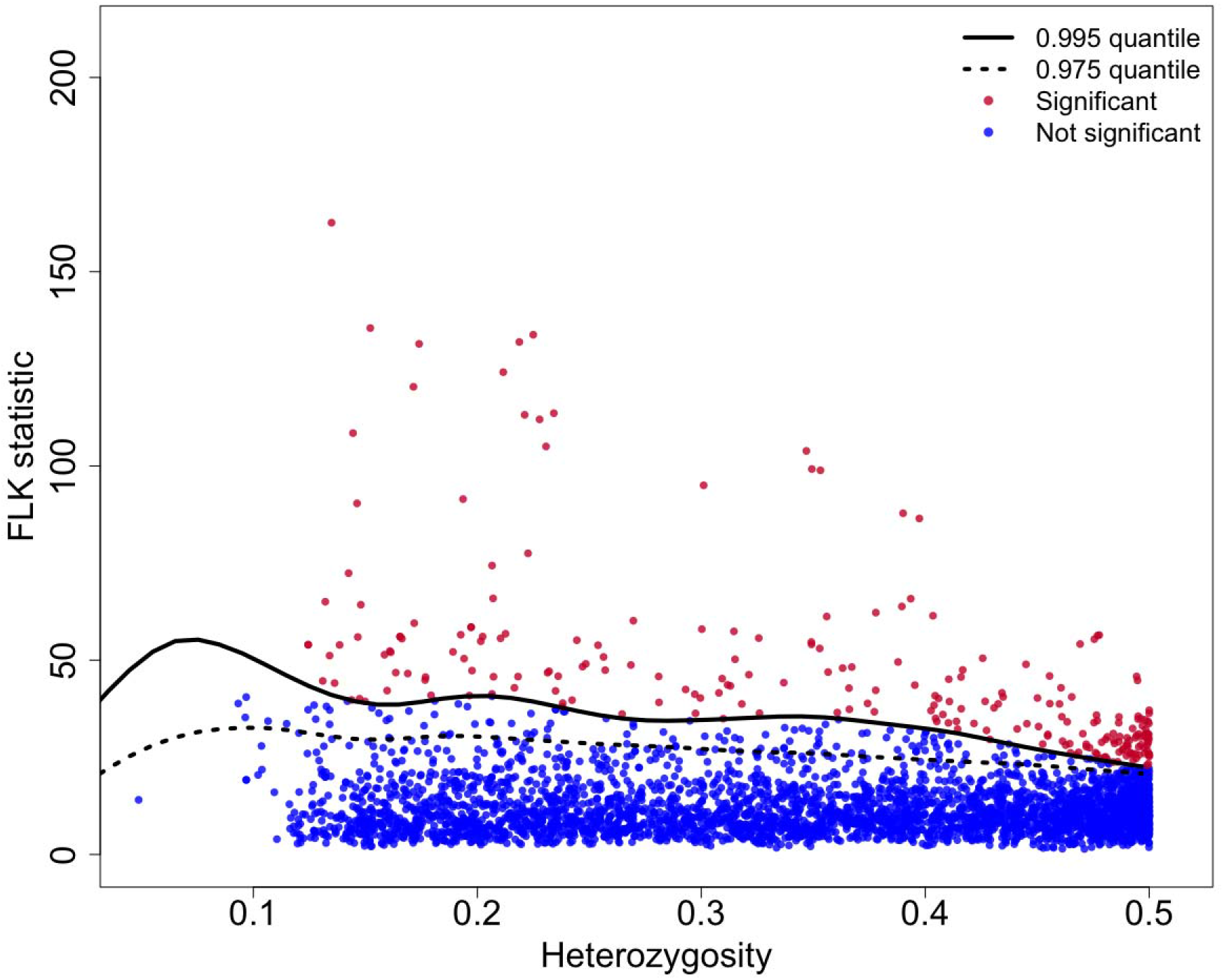
Genome-wide FLK test statistics and simulated 0.995 and 0.975 quantiles. The x-axis is the SNP heterozygosity, and the y-axis is the FLK statistic. The solid line indicates the significance threshold. Significant SNP outliers are highlighted with red color above the solid line, consistent with among-population differentiation above that expected from genetic drift alone.

### Genome-wide association analyses

Only one SNP (37855763-68-T/C, SNP ID assigned by DArT) was significantly associated with a trait (stipe length P-value = 2.03×10^−7^), with a minor allele frequency of 0.07 overall. The sequence containing SNP data is provided in Supplement Table 2.

### Detection of selection signatures

The FLK test detected potentially non-neutral outlier loci, which could have resulted from local adaptation involving linked sites. FLK statistics above the 0.995 quantiles were considered significant. A total of 249 SNPs were identified. The SNP with the highest FLK statistic (162.61) was 37850355-52-C/T. The heterozygosity of the SNP was 0.13.

## Discussion

In this study, we investigated the genetic population structure of wild sugar kelp (*Saccharina* sp.) populations in Northwest Atlantic using genome-wide SNP data. This structure is driven by mutation, drift, selection, and migration. The molecular methods used here can be applied to a variety of marine organisms to identify fine-scale population structure in nearshore coastal systems. Furthermore, we evaluated the phenotypic variation among wild samples. In general, the phenotypic patterns varied from one location to another which is expected given the variation in environmental conditions throughout the sampling region and the plasticity of kelp in response (Bartsch et al., 2008).

### Population structure

A clear population structure was observed among natural samples collected from the Northwest Atlantic. The sugar kelp from GOM and SNE formed three sub-populations, as demonstrated by analyses of pairwise F_ST_, PCoA, and admixture. The major genetic distinction between GOM and SNE suggests that Cape Cod acts as a biogeographic barrier for sugar kelp gene flow. Previous studies on larval transport reveal that species boundaries exist at Cape Cod largely set by the interaction of currents, depth distribution, and in the case of larvae, their pelagic duration (Pappalardo et al., 2015).

Overall, populations in the GOM showed three times more structure among locations than those found in SNE based on hierarchical AMOVA and pairwise F_ST_ (Table 2, Fig.4). The difference in the genetic population structure between the GOM and SNE was also qualitatively confirmed by PCoA (Fig. 5). The differentiation observed between *S. angustissima* (at Giant’s Staircase) and *S. latissima* was similar to that of *S. latissima* among locations (Fig. 5). The largest F_ST_ between *S. angustissima* and *S. latissima*, 0.29, was no larger than that of *S. latissima* samples between GOM and SNE (largest F_ST_ 0.32). Moreover, manipulative crosses between gametophytes obtained from *S. angustissima* and *S. latissima* can produce fertile hybrids (Umanzor et al., *In Prep*), indicating a lack of intrinsic reproductive isolating mechanisms. Nonetheless, previous ecological and molecular studies suggested divergence between these kelp forms, including habitat, morphology, and several organellar markers (Augyte et al., 2018). Common garden experiments further suggest a genetic basis driving the unique morphology of *S. angustissima* (Augyte at el., 2017; 2018).

The difference between the population structure of GOM and SNE might also be due to postglacial migration. Biogeographic reconstruction suggests the early expansion and diversification of complex kelp started in the northeast Pacific with dispersal and colonization to other regions via oceanographic flow through the Bering Strait and Arctic (Wares et al., 2001; Neiva et al., 2018; Starko et al., 2019). Divergence of western and eastern Atlantic populations predates the Last Glacial Maximum (Neiva et al., 2018) when an ice sheet extended onto the continental shelf and reached into the western Atlantic as far south as Long Island, New York (Maggs et al., 2008). Arctic populations with genetic affinities to the North Pacific and western Atlantic sugar kelp populations currently are hybridizing in parts of the Arctic, which may be typical of previous interglacial periods (Neiva et al., 2018). The southern range edge populations (SNE) analyzed here, in contrast, may remain more isolated and persist across climatic shifts or glacial vicariance events (Neiva et al., 2018).

### Candidate SNPs for morphology and selection signature

To our knowledge, no candidate genes have been reported for morphological traits in sugar kelp. In a closely related species *Saccharina japonica*, candidate genes have been reported for blade length and blade width (Wang et al., 2018). We found one SNP, 37855763-68-T/C, significantly associated with stipe length. More SNPs might have been discovered associated with other traits with a larger sample size. In the future, more samples should be collected for detecting the causal loci affecting sugar kelp morphology to better understand the genetic architecture of traits of economic interest.

There was no overlap between SNPs detected from GWAS and the FLK test because these tests exploit different signals in the data. As implemented here, the association analysis identified an SNP causing variation within locations, whereas the FLK test identified SNPs exceptionally divergent between locations. In addition, the association study adopted a stringent Bonferroni correction to eliminate false positives, resulting in a small set of significant SNPs. Finally, GWAS is a powerful statistical method for linking genotype to phenotype, but in the context of genomic variation caused by ongoing evolutionary forces such as selection, it is hampered by confounding selective pressures such as drift causing population structure (e.g., Liu et al., 2016).

### Genetic diversity within locations

Zhang et al (2015) reported using high-density SNP markers to assess kelp genetics for *Saccharina japonica*. To the best of our knowledge, our study is the first genetic study for *S. latissima* and putative species *S. angustissima* in the Northwest Atlantic region using genomewide SNP data. Previously, genetic diversity in eastern Maine was assessed using 12 microsatellite markers, and samples from five intertidal locations (i.e., Penobscot Bay, Frenchman Bay, Cobscook Bay, Englishman Bay, and Starboard Cove) (Breton et al., 2018). Their range of observed heterozygosity was estimated from 0.283 to 0.339. In European sugar kelp populations, genetic diversity has been estimated with the same 12 microsatellite markers and reported much higher observed heterozygosity (Nielsen et al., 2016; Paulino et al., 2016). For example, Møller Nielsen et al., (2016) reported that the observed heterozygosity was 0.560. Lower genetic diversity in western North Atlantic benthic taxa relative to Europe has been reported for multiple taxa (Ware and Cunningham 2001) and hypothesized to result from postglacial recolonization from East to West. The purpose of the conStruct model is to raise hypotheses concerning the population history of a species (Bradburd et al. 2018). In the context of these previous observations, we propose two of the layers we found to be distinct colonization events from east to west in the Atlantic, while the third layer derives from the separation of subpopulations due to the Cape Code barrier. Admixture and divergence, we believe, could be the odd geographic pattern of genetic differentiation observed.

Compared with GOM (mean observed heterozygosity = 0.29), SNE exhibited modestly lower (p-value < 0.05) genetic diversity (mean observed heterozygosity = 0.27). This difference could be due to either a lower effective population size in SNE populations at the range limit, or that greater migration to GOM than to SNE occurred during the second re-colonization event. In the future, genome-wide SNP data should be applied to sugar kelp genetic diversity globally. Such information can show whether the SNE population exhibits adaptation to higher water temperatures or is merely bottlenecked. That inference will be important for improving our understanding of how kelp responds to climate change (Hoffman and Sgrò 2011).

## Conclusion

In the Northwest Atlantic, the coastal geography, currents, vicariance events, and genetic drift have led to ecological diversification and separation of *Saccharina latissima* gene pools in the GOM and SNE subpopulations. Cape Cod was confirmed to be a longstanding biogeographic barrier between GOM and SNE sugar kelp, but perhaps not completely impermeable in the recent past. It is notable that despite this deep regional population structure, the genetic variation found within locations accounted for the greatest proportion of the total, indicating abundant local standing diversity for selection to act on. Breeding among samples between these two regions is currently managed and prohibited due to conservation concerns. Our results indicate that these regions might have already exchanged migrants on their own over time. Nonetheless, as in other similar studies including Evankow et al., (2019), our recommendation supports the precautionary principle to only use regional ecotypes for cultivation and to not transport kelp strains across different regions (e.g. between GOM and SNE, in this case). We also reveal a novel SNP association with a morphological trait and additional candidates for selection in sugar kelp. Our study can be used to guide conservation and management decisions as well as future kelp breeding research.

## Acknowledgments

Funding support from the U.S. Dept. of Energy ARPA-E (DE-AR0000915). We acknowledge Christina Marie Rochus for suggestions on the data analyses.

## Author contribution statement

XM and SA wrote the manuscript draft. XM and MH performed the analysis and discussed results together. XM, SA, and MH revised the manuscript together. MPH guided population genetic analyses and edited the manuscript. CY offered guidance on collections and DB, SL and SA collected samples. DB, SL, SA, SU and MM-R phenotyped the samples. KRR guided analyses, provided computational resources and edited the manuscript; SL, CY and J-LJ led the project and edited the manuscript.

## Data Availability

Supplemental data is available online

