## Supplemental Figure 1 for "Population genetics of sugar kelp in the Northwest Atlantic region using genome-wide markers": Sugar kelp_Supplemental Figure 1.docx

Visualization of layers (K) contributions from conStruct results using MCMC=50,000 iterations with a burn-in of 500 iterations that were discarded. The layer contribution is estimated in the software as proportion of variance being explained by adding that layer to the total variance. Results shown for a) all samples with K=1 to 6; b) all samples with K=3 and K=4; c) samples within GOM from K=1 to 4;

a

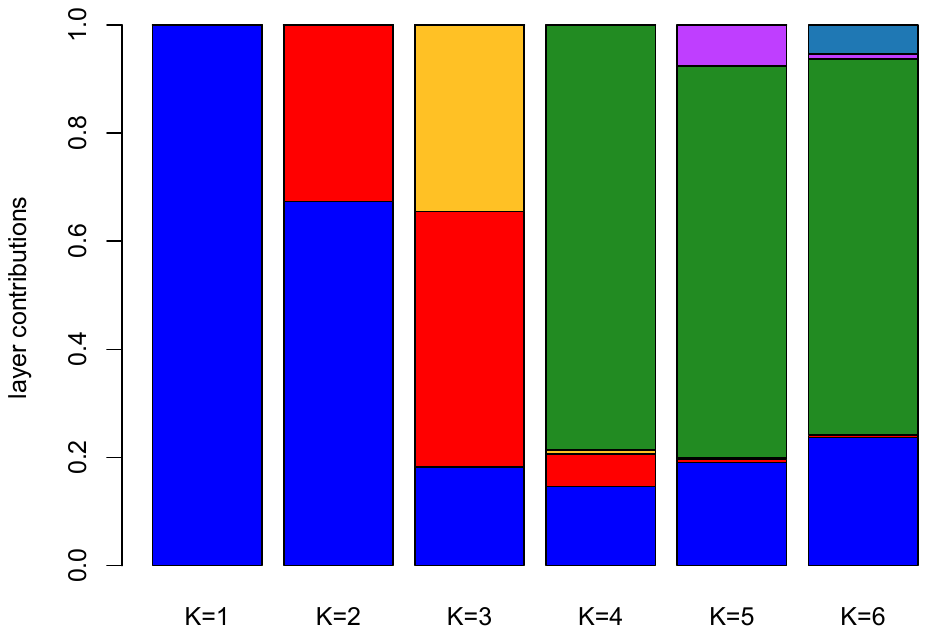

b.

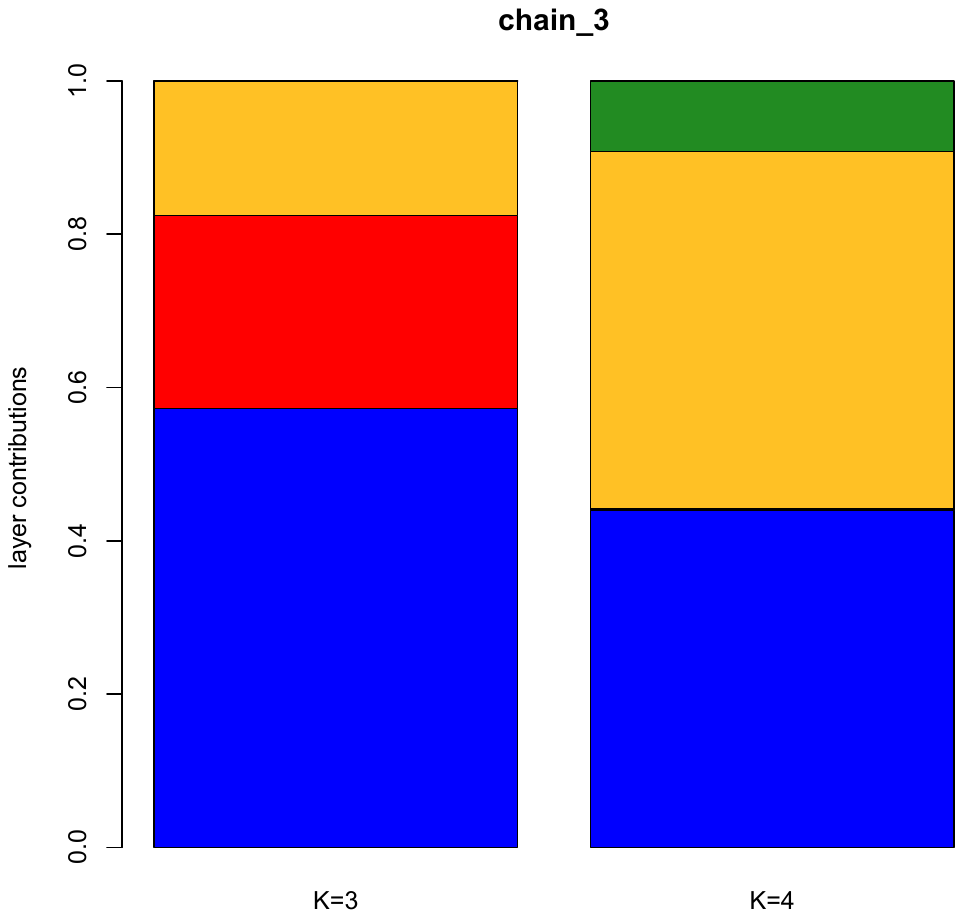

c.

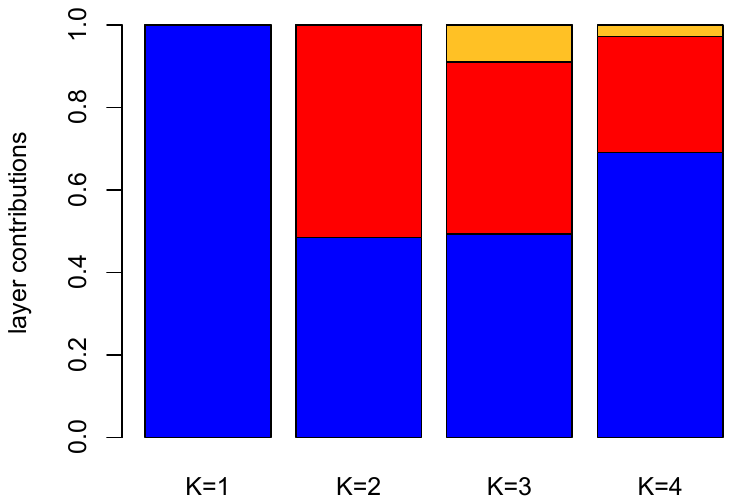
