## Supplemental Table 1 for "Population genetics of sugar kelp in the Northwest Atlantic region using genome-wide markers": Sugar_kelp_Supplement Table 1_Fst.docx

Supplement Table 1. Pair-wise F_ST_ values across locations.

|  | Fort Wetherill | Giant's Staircase | Cape Cod Canal | Downeast Institute | Fisher's Island | Fort Stark | Isles of Shoals | Newcastle | Black Ledge | Lubec Dock | Lubec Light | Mt Desert Rock | Nubble Light | Orr's Island | Pine Island |
| --- | --- | --- | --- | --- | --- | --- | --- | --- | --- | --- | --- | --- | --- | --- | --- |
| Fort Wetherill | NA | NA | NA | NA | NA | NA | NA | NA | NA | NA | NA | NA | NA | NA | NA |
| Giant's Staircase | 0.27184 | NA | NA | NA | NA | NA | NA | NA | NA | NA | NA | NA | NA | NA | NA |
| Cape Cod Canal | 0.22431 | 0.16681 | NA | NA | NA | NA | NA | NA | NA | NA | NA | NA | NA | NA | NA |
| Downeast Institute | 0.25854 | 0.18600 | 0.10880 | NA | NA | NA | NA | NA | NA | NA | NA | NA | NA | NA | NA |
| Fisher's Island | 0.04655 | 0.28518 | 0.23878 | 0.27415 | NA | NA | NA | NA | NA | NA | NA | NA | NA | NA | NA |
| Fort Stark | 0.23719 | 0.13625 | 0.08636 | 0.10241 | 0.25012 | NA | NA | NA | NA | NA | NA | NA | NA | NA | NA |
| Isles of Shoals | 0.26134 | 0.17397 | 0.07705 | 0.10555 | 0.27549 | 0.05616 | NA | NA | NA | NA | NA | NA | NA | NA | NA |
| Newcastle | 0.23608 | 0.14153 | 0.08529 | 0.09765 | 0.25018 | 0.00641 | 0.05188 | NA | NA | NA | NA | NA | NA | NA | NA |
| Black Ledge | 0.05388 | 0.29415 | 0.23798 | 0.27036 | 0.00992 | 0.25226 | 0.27145 | 0.25135 | NA | NA | NA | NA | NA | NA | NA |
| Lubec Dock | 0.28490 | 0.21217 | 0.15298 | 0.10653 | 0.29520 | 0.14964 | 0.15686 | 0.14177 | 0.29701 | NA | NA | NA | NA | NA | NA |
| Lubec Light | 0.28434 | 0.21743 | 0.15118 | 0.10344 | 0.29661 | 0.14544 | 0.15658 | 0.13954 | 0.29540 | 0.00546 | NA | NA | NA | NA | NA |
| Mt Desert Rock | 0.29406 | 0.20581 | 0.20879 | 0.18633 | 0.30988 | 0.16052 | 0.20120 | 0.16569 | 0.31714 | 0.22287 | 0.22703 | NA | NA | NA | NA |
| Nubble Light | 0.25074 | 0.13817 | 0.11353 | 0.12641 | 0.26338 | 0.02299 | 0.07901 | 0.03079 | 0.27041 | 0.16970 | 0.17148 | 0.16452 | NA | NA | NA |
| Orr's Island | 0.22505 | 0.12187 | 0.08845 | 0.09070 | 0.23553 | 0.08500 | 0.09470 | 0.08071 | 0.23710 | 0.12798 | 0.13194 | 0.19369 | 0.11676 | NA | NA |
| Pine Island | 0.03784 | 0.27830 | 0.21834 | 0.24897 | 0.00277 | 0.22617 | 0.25301 | 0.22518 | 0.00875 | 0.28091 | 0.27897 | 0.30262 | 0.24776 | 0.21065 | NA |
