## Supplemental Table 2 for "Population genetics of sugar kelp in the Northwest Atlantic region using genome-wide markers": Sugar_kelp_Supplemental Table 2.docx

Supplemental Table 2. Reference and alternative sequence reads for significant single nucleotide markers (SNPs) in the Genome Wide Association analysis (GWAS) and top ten significant SNPs from extended Lewontin and Krakauer (FLK) selection signature test

| Statistical test | Significant SNPs | Reference read |
| --- | --- | --- |
| GWAS | 37855763-68-T/C | TGCAGTATTCTCCCCCCTCTGGTGTAGGACATGGGTGGCCAAAGAGGGTCAAGCCTGGGCACTGGGCAT |
| FLK | 37850355-52-C/T | TGCAGGGCCCTTGCCGCCCTCTGAGGGTACGGCTAAAACCAAAACGAACTTTCCCCCCCG |
|  | 37832954-53-G/A | TGCAGGCATAGGTTAAGAGTGTGCCAGCACCATTGTCTTGTGCATCAACATGAGCCCCGTGACGAATGA |
|  | 37862869-67-C/T | TGCAGTAGCAGCCAAGCCTGCTACTTGCGCCTACGGGACTCTGACCACTTCACTTAACTCTTGAAAACC |
|  | 37841131-41-C/T | TGCAGCAACCCCAGCAGCAGCAACAGCAACAGCAGCAGCAACAGCAGCAGCAACATCACCGCCACCGAC |
|  | 37857606-5-T/C | TGCAGTAGCAACGATAGATAGACTAGTTGTGAGAAACGCAGTCGGGTTGCGGACGTTCGTCGAGTCGGG |
|  | 37848571-13-A/G | TGCAGGACGCCATAGGCTGGTGGCAGCGGGAGAGCTCGCGGTTCGACGCCAGAGTGTGGACGGGGGAAC |
|  | 37853585-39-G/A | TGCAGAGTAAGACCATGAGGGTTGATTGTAGCAAGGGGCGTCCTCACCG |
|  | 37857066-8-G/A | TGCAGTAAGCGAACGATTAAACAAATCGATGGTACCTGTGCATATGGTAACTATAGGTATCACGAGTAT |
|  | 37854090-21-C/T | TGCAGGTGTGGTGACGGCCGACGGGCTCTCTCGCGTGAACATCAGTGGCAACACCTGGTTTCACAACAA |
|  | 37847908-28-G/A | TGCAGCACGCCGACCGCAGGCAACTCGAGACGACTTGCGCCGAATTTAGTCCAACGCGCTACGAGCGCA |
|  |  | Alternative read |
| GWAS | 37855763-68-T/C | TGCAGTATTCTCCCCCCTCTGGTGTAGGACATGGGTGGCCAAAGAGGGTCAAGCCTGGGCACTGGGCAC |
| FLK | 37850355-52-C/T | TGCAGGGCCCTTGCCGCCCTCTGAGGGTACGGCTAAAACCAAAACGAACTTTTCCCCCCG |
|  | 37832954-53-G/A | TGCAGGCATAGGTTAAGAGTGTGCCAGCACCATTGTCTTGTGCATCAACATGAACCCCGTGACGAATGA |
|  | 37862869-67-C/T | TGCAGTAGCAGCCAAGCCTGCTACTTGCGCCTACGGGACTCTGACCACTTCACTTAACTCTTGAAAATC |
|  | 37841131-41-C/T | TGCAGCAACCCCAGCATCAGCAACAGCAACAGCAGCAGCAACAGCAGCAGCAACATCACCGCCACCGAC |
|  | 37857606-5-T/C | TGCAGCAGCAACGATAGATAGACTAGTTGTGAGAAACGCAGTCGGGTTGCGGACGTTCGTCGAGTCGGG |
|  | 37848571-13-A/G | TGCAGGACGCCATGGGCTGGTGGCAGCGGGAGAGCTCGCGGTTCGACGCCAGAGTGTGGACGGGGGAAC |
|  | 37853585-39-G/A | ATGCAGAGTAAGACCATGAGGGTTGATTGTAGCAAGGGGCATCCTCACCG |
|  | 37857066-8-G/A | TGCAGTAAGCGAACGATTAAACAAATCGATGGTACCTGTGAATATGGTAACTATAGGTATCACGAGTAT |
|  | 37854090-21-C/T | TGCAGGTGTGGTGACGGCCGATAGGCTCTCTCGCGTGAACATCAGTGGCAACACCTGGTTTCACAACAA |
|  | 37847908-28-G/A | TGCAGCACGCCGACCGCAGGCAACTCGAAACGACTTGCGCCGAATTTAGTCCAACGCGCTACGAGCGCA |
